# Population genomics indicate three different modes of divergence and speciation with gene flow in the green-winged teal duck complex

**DOI:** 10.1101/2021.09.10.459850

**Authors:** Fern Spaulding, Jessica F. McLaughlin, Kevin G. McCracken, Travis C. Glenn, Kevin Winker

**Affiliations:** University of Alaska Museum, University of Alaska Fairbanks, Fairbanks, AK, USA; Department of Biology and Wildlife, University of Alaska Fairbanks, USA; Sam Noble Oklahoma Museum of Natural History, University of Oklahoma, Norman, OK, USA; Department of Biology, University of Miami, Coral Gables, FL, USA; Department of Environmental Health Science, University of Georgia, Athens, GA, USA

**Keywords:** Anatidae, waterfowl phylogenetics, Beringia, speciation with gene flow, parapatric speciation, allopatric speciation

## Abstract

The processes leading to divergence and speciation can differ broadly among taxa with different life histories. We examine these processes in a small clade of ducks with historically uncertain relationships and species limits. The green-winged teal (*Anas crecca*) complex is a Holarctic species of dabbling duck currently categorized as three subspecies (*Anas crecca crecca*, *A. c. nimia*, and *A. c. carolinensis*) with a close relative, the yellow-billed teal (*Anas flavirostris*) from South America. We examined divergence and speciation patterns in this group, determining their phylogenetic relationships and the presence and levels of gene flow among lineages using both mitochondrial and genome-wide nuclear DNA obtained from 1,393 ultraconserved element (UCE) loci. Phylogenetic relationships using nuclear DNA among these taxa showed *A. c. crecca*, *A. c. nimia*, and *A. c. carolinensis* clustering together to form one polytomous clade, with *A.flavirostris* sister to this clade. This relationship can be summarized as (*crecca*, *nimia*, *carolinensis*)(*flavirostris*). However, whole mitogenomes revealed a different phylogeny: (*crecca*, *nimia)(carolinensis*, *flavirostris*). The best demographic model for key pairwise comparisons supported divergence with gene flow as the probable speciation mechanism in all three contrasts (*crecca*−*nimia*, *crecca*−*carolinensis*, and *carolinensis*−*flavirostris*). Given prior work, gene flow was expected among the Holarctic taxa, but gene flow between North American *carolinensis* and South American *flavirostris* (*M* ∼0.1 - 0.4 individuals/generation), albeit low, was not expected. Three geographically oriented modes of divergence are likely involved in the diversification of this complex: heteropatric (*crecca*−*nimia*), parapatric (*crecca*−*carolinensis*), and (mostly) allopatric (*carolinensis*−*flavirostris*). Ultraconserved elements are a powerful tool for simultaneously studying systematics and population genomics in systems like this.

**Graphical Abstract:** 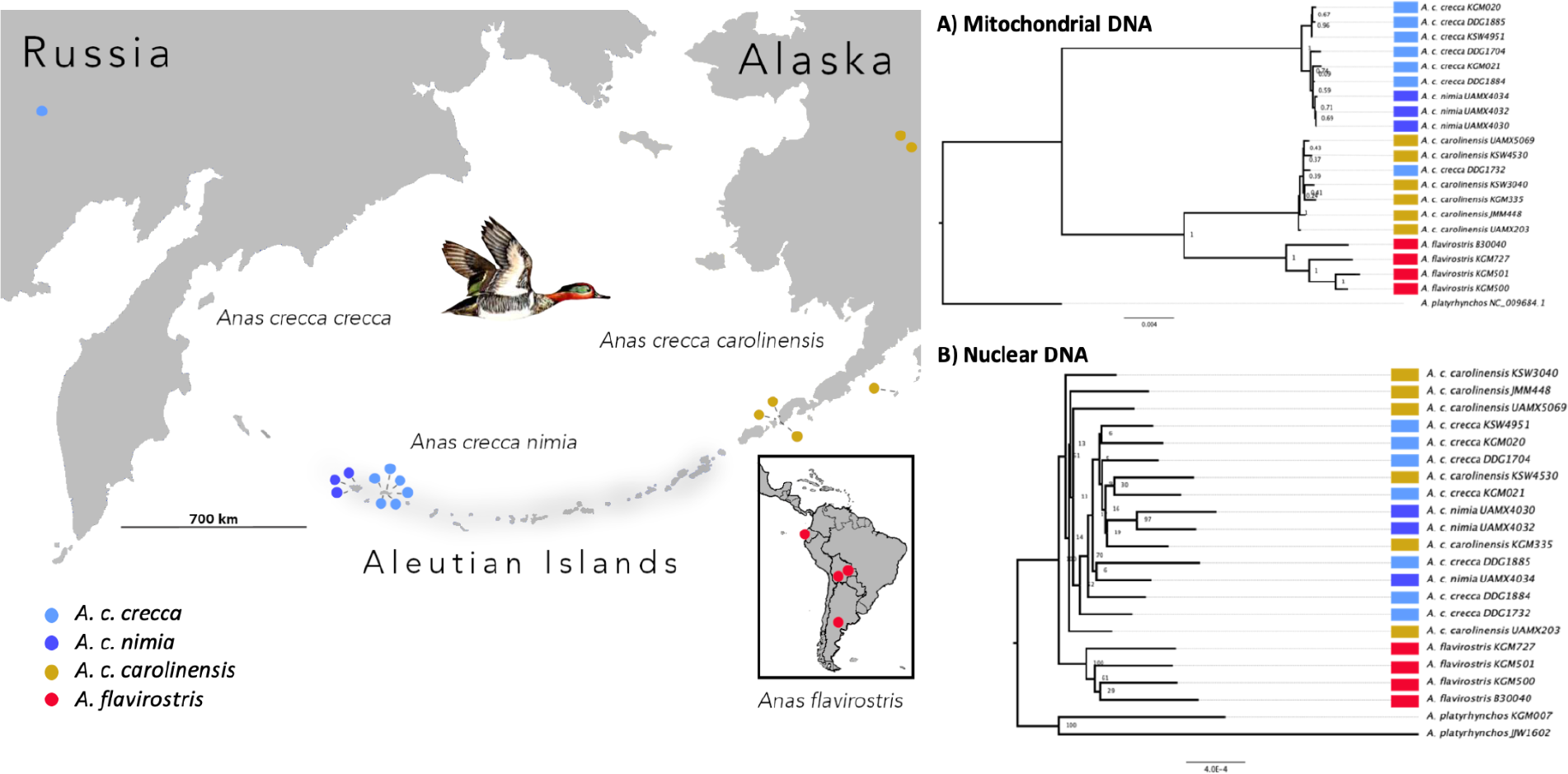

## 1. Introduction

A fundamental goal of evolutionary biology is to understand different modes of divergence and speciation causing the generation of biodiversity. Speciation is the process of evolutionary divergence that often results in distinct groups of organisms recognized as species, which are essentially reproductively isolated from each other (Mayr, 1942; Price, 2008). This process has historically been categorized in three main modes: sympatric, parapatric, and allopatric, each of which is based on spatial relationships between the diverging populations (Fitzpatrick et al., 2008). Classic allopatric speciation is the divergence of populations that are geographically separated from each other with no movement of individuals between them (Mayr, 1942; Gavrilets, 2003, 2004; Coyne & Orr, 2004; Fitzpatrick et al., 2008; Price, 2008).

Sympatric speciation involves the evolution of reproductive isolation while the ranges of populations overlap (Coyne & Orr, 2004). Parapatric speciation is the origin of new species in which gene flow occurs across a spatially restricted contact zone such that only a fraction of each population has a high probability of emigrating or of interacting with immigrants (Smith, 1955; Endler, 1977; Futuyma & Mayer, 1980, Fitzpatrick et al., 2008). These geographic modes of speciation can be considered to exist on a continuum, on which allopatric and sympatric speciation represent the endpoints of different amounts of gene flow (zero to maximum), whereas parapatric speciation occupies the space representing intermediate gene flow between these extremes (Butlin et al., 2008; Gavrilets, 2014). In speciation theory, if the connectivity between two populations is not broken, and gene flow persists, then parapatric speciation models apply (speciation with gene flow, in a non-sympatric distribution; Gavrilets, 2004). The frequency of parapatric speciation in nature is uncertain (Coyne & Orr, 2004; Price, 2008), and it has been a relatively neglected area in speciation research (Gavrilets, 2004).

The study of speciation has often focused on these geographic modes of divergence.

There are major debates about whether allopatric speciation is predominant (e.g. Coyne & Orr, 2004), whether sympatric speciation is possible, and if so how frequently it might occur (e.g. Berlocher & Feder, 2002; Bolnick & Fitzpatrick, 2007). In some cases, geographic isolation alone has been shown to drive divergence when isolated populations reside in allopatry. For example, allopatric speciation is common in molluscs (78% of species pairs), driven in part by their low dispersal rates (Hernández-Hernández et al., 2021). Similarly, among terrestrial snails that live on island archipelagos, allopatric species pairs predominate (96%), with most species being endemic to a single island (Holland & Cowie, 2009; Rundell, 2008; Jordaens et al., 2009; Hernández-Hernández et al., 2021).

Though the geographic distribution of populations is important for understanding speciation, other factors such as ecological, environmental, and behavioral differences between populations are increasingly seen as important drivers of population divergence and speciation (Schluter, 2001; McKinnon et al., 2004; Ruegg et al., 2012; Verzijden et al., 2012, Withrow et al., 2014). While these factors can operate within each category of the geographically oriented modes of speciation (allopatric, parapatric, and sympatric; Gavrilets, 2003), their presence is particularly important for enabling parapatric or sympatric speciation to progress (speciation with gene flow; Rundle & Nosil, 2005; Nosil, 2008; Feder et al., 2012). For example, ecological and sexual selection are considered major evolutionary forces that often drive insect speciation (Arnqvist et al., 2000; Forbes et al., 2017; Hernández-Hernández et al., 2021), and spatial relationships can become less important. A classic example of this occurs in apple maggot flies (*Rhagoletis pomonella*), in which ecological divergence enables speciation through divergent adaptation to the preferred host plants of these phytophagous flies (Filchak et al., 2000). Another example of ecological divergence occurs in *Timema* walking stick insects where ecotypes live on different host plants (Nosil et al., 2008), in which greater reproductive isolation evolves between populations adapting to contrasting environments than between populations adapting to similar environments (Rice & Hostert, 1993; Schluter & Nagel, 1995; Schluter, 2009). Environmental factors are also often an important driver of divergence (Hernández-Hernández et al., 2021), and this occurs in many vertebrate groups, including salamanders (Kozak & Wiens, 2010), frogs (Moen & Wiens, 2017), birds (Cooney et al., 2016), and mammals (Castro-Insua et al., 2018).

Finally, behavior is commonly involved in prezygotic isolating barriers, including ecological and behavioral differences between species (Hernández-Hernández et al., 2021). For example, species-specific vocalization and communication are often important reproductive isolating mechanisms in birds and frogs (e.g., Edwards et al., 2005; Hoskin et al., 2005; Boul et al., 2007; Uy et al., 2018).

In birds, allopatric speciation has historically been thought to be the main route to speciation (Mayr, 1963; Coyne & Orr, 2004; Price, 2008). However, genomic data increasingly identify groups that do not fit this model and instead indicate that speciation has progressed with at least some gene flow (Mallet et al., 2016; Morales et al., 2017; Peñalba et al., 2019; Rheindt & Edwards, 2011; Zarza et al., 2016, McLaughlin et al., 2020; Winker, 2021). One example of how the allopatric speciation model is not an ideal fit for speciation in birds is that many long-distance seasonally migratory birds often exhibit semiannual transcontinental and transoceanic movements that can prevent diverging populations from undergoing long periods of strict allopatry (Winker, 2010; Peters et al., 2012), increasing the likelihood of divergence with gene flow.

Holarctic avian taxa breeding across the Northern Hemisphere provide an excellent opportunity to study possible speciation with gene flow, given that many of these birds are long-distance migrants that seasonally migrate to the Northern Hemisphere to breed. Here we focus on divergence in an avian taxonomic complex in Beringia, a geographic region that extends from the Russian Far East across Alaska and into western Canada and which has experienced dynamic fluctuations in climate throughout the Pleistocene. While much of the Northern Hemisphere was covered by ice sheets during past glaciations, the lowlands of Beringia remained free of ice, providing a refuge for high-latitude flora and fauna (Elias & Brigham-Grette, 2013). These cyclic fluctuations in climate have had genetic consequences on Beringian taxa (Hewitt, 1996), for example creating diverse patterns of differentiation in Holarctic birds (e.g. Zink et al., 1995; Drovetski et al., 2004; Buehler & Baker, 2005; Humphries & Winker, 2011; Peters et al., 2014). Previous work on speciation and divergence in birds across Beringia has shown that speciation with gene flow is common (Winker et al., 2013, 2018; Peters et al., 2014; McLaughlin et al., 2020). In the green-winged teal (*Anas crecca* and its subspecies), cyclic climatic fluctuations seem to have caused incomplete parapatric speciation (divergence with gene flow) between Asian and North American populations (*A. c. crecca* and *carolinensis*; Peters et al., 2012). A sedentary population in the Aleutian Islands (*A. c. nimia*) is also undergoing divergence with gene flow from the migratory Asian mainland subspecies (*A. c. crecca*), which performs seasonal migrations through the range of *nimia* in a heteropatric (seasonally sympatric) geographic relationship (Winker et al., 2013). The closest relative of these lineages is the South American yellow-billed teal (*A. flavirostris*), whose exact relationship with *A. crecca* (*sensu lato*) remains unresolved (Peters et al., 2012).

In this study, we examined the green-winged teal complex comprising three subspecies of green-winged teal in the Northern Hemisphere (the migratory Eurasian common teal (*Anas crecca crecca*) and North American green-winged teal (*A. c. carolinensis*), and the sedentary Aleutian green-winged teal (*A. c. nimia*)), and a closely related sister species, the yellow-billed teal (*A. flavirostris*) in the Southern Hemisphere (Fig. 1). This latter South American species is known to be closely related to *A. crecca*, but it has previously demonstrated conflicting relationships between mitochondrial and nuclear phylogenies, and it seems likely to represent a classic case of allopatric speciation (Peters et al., 2012). Taxonomically, the current species tree is (*A. c. crecca*, *A. c. nimia*, *A. c. carolinensis*)(*A. flavirostris*). However, mitochondrially the phylogeny is (*A. c. crecca*, *A. c. nimia*)(*A. c. carolinensis*, *A. flavirostris)* (Johnson & Sorenson, 1999; Gonzalez et al., 2009). Here we bring a genomic-scale dataset to this system, asking first what are the phylogenetic relationships in the group? Second, how much gene flow has occurred? And, finally, what modes of divergence and speciation are prevalent in this group?

**Figure 1.**
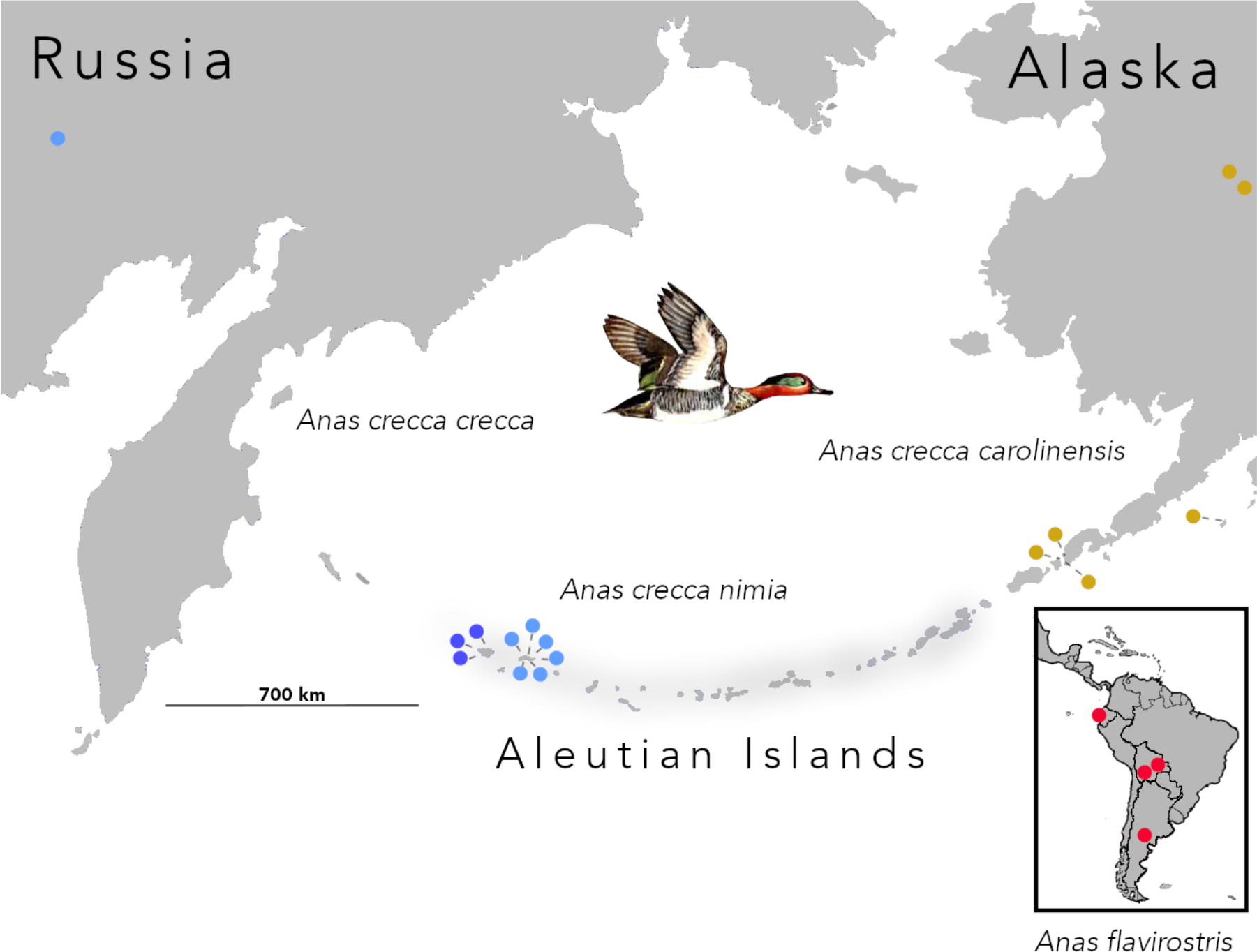
Distribution of samples used in this study, focusing on Beringia and South America (inset). *A. c. crecca* occurs throughout most of Eurasia and seasonally migrates through the western Aleutians (light blue; 7 individuals), *A. c. nimia* occurs as a resident in the Aleutian Islands (dark blue; 3 individuals), and *A. c. carolinensis* occurs across North America (gold; 6 individuals). The yellow-billed teal, *Anas flavirostris* (red; 4 individuals) is a South American sister taxon. See Supplementary Table S1 for specimen details. Illustration of green-winged teal courtesy of USFWS (Hines, 1963).

## 2. Materials and methods

### 2.1. Study design

We used ultraconserved elements (UCEs) from the nuclear genome as our primary genetic markers. UCEs allow us to examine thousands of orthologous loci, providing insight into the divergence and speciation processes from populations to deeper relationships (Faircloth et al., 2012; Everson et al., 2019). In addition to obtaining thousands of nuclear loci, a positive byproduct of producing UCE data is that high-quality complete mitogenomes are also generated for each individual. Pairing UCEs with the complete mitogenomes for each individual allowed us to compare the phylogenetic relationships between nuclear and mitogenomic data. Given that there is often a lack of concordance between mitochondrial and nuclear DNA (mtDNA and nuDNA) regarding respective estimates of divergence (Humphries & Winker, 2011; Peters et al., 2014), using both types of molecular markers can improve our understanding of lineage relationships and species limits (Rubinoff & Holland, 2005; Edwards & Bensch, 2009; Humphries & Winker, 2011). Using both mtDNA and nuDNA simultaneously can also resolve potential discrepancies between previously published phylogenies.

### 2.2. Sampling, DNA extraction, and sequencing

We sampled the following numbers of individuals from each taxon: 7 *Anas c. crecca*, 6 *A. c. carolinensis*, 3 *A. c. nimia*, 4 *A. flavirostris* (Fig. 1), and 2 *A. platyrhynchos* were used as an outgroup. In this study, we consider each named taxon a population, so we are conducting comparisons at both the subspecies and species level. Theory considers an optimal sample size for coalescent-based genomic analyses to be 8 individuals (or haplotypes) per population when estimating population parameters (e.g., θ = 4*Ne*μ; Felsenstein, 2005). However, it has been demonstrated that key demographic parameter estimates in lineages with divergences of these depths (i.e., subspecies and species) are generally resilient to lower sample sizes (McLaughlin & Winker, 2020). It has also been shown that, in general, relatively small sample sizes are sufficient when estimating interpopulation divergence and genetic diversity when using thousands of loci (Nazareno et al., 2017; McLaughlin & Winker, 2020). In our population-level analyses, each allele is called, which effectively doubles the sample size of each taxon (two haplotypes per individual). A minimum reliable sample size varies from taxon to taxon (though is frequently much lower than 8), and is dependent on the demographic parameters of interest and the divergence levels between the taxa (McLaughlin & Winker, 2020). Here we are particularly interested in the possible presence and levels of gene flow (*m*), which when using our methods of analysis appear to be relatively consistent even when sample sizes are small (McLaughlin & Winker, 2020). For phylogenetic systematics analyses we used a single consensus sequence per individual (Faircloth, 2016). Each sample was obtained from high-quality, vouchered tissue samples from wild individuals (Table S1). DNA extractions followed standard protocol for animal tissues using the QIAGEN DNeasy Blood + Tissue Extraction Kit (QIAGEN, 2006).

Our sequencing procedures followed Glenn et al. (2019). In short, we prepared dual-indexed DNA libraries which were quantified using a Qubit fluorimeter (Invitrogen, Inc., Carlsbad, CA, USA). We then enriched the samples with 5,060 UCE loci using the Tetrapods-UCE-5Kv1 kit from MYcroarray following version 1.5 of the UCE enrichment protocol and version 2.4 of the post-enrichment amplification protocol (http://ultraconserved.org). The resulting pool was then sequenced using a paired-end 150 bp (PE150) protocol on an Illumina HiSeq 2500 using three lanes (Illumina, Inc., San Diego, CA, USA; UCLA Neuroscience Genomics Core).

### 2.3. Bioinformatics and UCE pipeline

Our bioinformatics pipeline followed that of Winker et al. (2018). Briefly, raw and untrimmed FASTQ sequence data that contained low-quality bases were removed using Illumiprocessor (v.2.0.6; Faircloth, 2013), which incorporates Trimmomatic (v.0.32-1; Bolger et al., 2014). Our next steps used the package PHYLUCE (v.1.5.0; Faircloth, 2016), which identifies conserved orthologous loci that are then used as our reference set of UCE loci to call variants in the focal, ingroup individuals. We used the mallard (*Anas platyrhynchos*) to build a UCE reference. We first combined sequence read files from two individuals into two read files (Table S1). We assembled these reads de novo using Trinity (v.2.4.0; Grabherr et al., 2011) on Galaxy (v.2.4; Afgan et al., 2016), then found and extracted UCE loci using PHYLUCE by matching the contigs to the probe set used. The resulting sequences were saved as a reference FASTA file. Next, for each ingroup individual, we combined singletons (reads that lost their pair) with read1 files. The PHYLUCE dependencies such as BWA-MEM (v.0.7.7; Li & Durbin, 2009; Li, 2013), SAMtools (v.0.1.19; Li et al., 2009), and PICARD (v.1.106; http://broadinstitute.github.io/picard) were used to index the reference sequence and align the unassembled raw reads of each individual against the mallard UCE reference using default parameters. We followed the population genomics pipeline developed by Harvey et al. (2016), which includes using Genome Analysis Toolkit (GATK, v.3.3.0; McKenna et al., 2010) to call and restrict data to high-quality SNPs (Q30).

VCFtools (v.0.1.13; Danecek et al., 2011) was used to cull sites without complete data to filter the high-quality SNPs to create a complete matrix (all individuals represented at all loci) with a minimum genotype quality (Phred) score of 10.0 (which equates to 90% confidence). Our final FASTA file contained only high-quality, confidently sequenced loci, producing a 100% complete data matrix for analyses. The high-quality, complete-data VCF file was thinned to 1 SNP/locus using VCFtools and made biallelic by filtering out loci that had more than two alleles. The thinned, biallelic VCF file then was used in our demographic model analysis. We used BLASTn on NCBI to identify sex linked loci (Z-linked) using our high-quality FASTA data. We used the mallard genome (*Anas platyrhynchos*, IASCAAS_PekingDuck_PBH1.5) to identify hits for Z-linked loci, which were removed from our thinned biallelic VCF using the script find_chrom.py (v.1.2; https://github.com/jfmclaughlin92/beringia_scripts).

The biallelic VCF dataset was converted using PGDSpider (v.2.0.9.1; Lischer & Excoffier, 2012) into the appropriate format for analysis using the R package adegenet (v.2.0.1; Jombart & Ahmed, 2011). Adegenet was used to calculate Hardy-Weinberg equilibrium, observed (*Ho*) and expected heterozygosities (*He*) , population level *FST* values, principal component analysis (PCA), and perform assignment tests using discriminant analysis of principal components (DAPC). A paired *t*-test was used to test for differences between expected and observed heterozygosity (Jombart & Ahmed, 2011).

Although we tested demographic models of divergence and gene flow, Hardy-Weinberg equilibrium calculations offer an independent assessment of gene flow because unequal values for observed and expected heterozygosity suggest that one of the assumptions of Hardy-Weinberg equilibrium is not met (i.e., absence gene flow). To calculate nucleotide diversity (π), we created a concatenated FASTA of all individuals using catfasta2phyml.pl (https://github.com/nylander/catfasta2phyml). The resulting file was then imported into MEGA (v.10; Kumar et al., 2018), and nucleotide diversity was calculated using the maximum composite likelihood method.

### 2.4. Mitogenomic pipeline

Mitogenomic sequence data were obtained as a byproduct from UCE sequencing. The backbone of our mitogenomic analysis is similar to that of our UCE dataset and follows many of the same steps. Our pipeline again centered on PHYLUCE and followed the mitogenomic pipeline used by Everson et al. (2019). Briefly, we used the complete mitogenome of the mallard (*Anas platyrhynchos,* NC_009684.1) as a reference. The read1 and read2 files were mapped to the reference and indexed using BWA and SAMtools. This was followed by using PICARD to clean the alignments, add read group header information, and remove PCR and sequencing duplicates. Using the GATK module UnifiedGenotyper, SNPs were called for each individual against the reference, and GATK was used to call and realign around indels, call and annotate SNPs, and filter SNPs around indels using the IndelRealigner module, which incorporates the merged interval output created by the RealignerTargetCreator module. We restricted the data to high-quality SNPs by adding a quality filter (Q30) before converting the resulting VCF file to a FASTA file using GATK. Mitochondrial divergence estimates were also calculated using whole mitogenomes to enable comparison with previous studies (Johnson & Sorenson, 1999; Humphries & Winker, 2011). Divergence estimates were calculated in MEGA using the between-group mean distance option.

### 2.5. Phylogenetic analysis

We used the PHYLUCE phylogenetic systematics pipeline (Faircloth, 2016) to generate a single UCE consensus sequence per locus/individual (heterozygotes are coded using IUPAC codes). We next used RAxML (using raxmlGUI v.1.5; Silvestro & Michalak, 2012) to reconstruct a maximum likelihood phylogeny. We used ModelTest-NG (v.0.1.7; Darriba et al., 2020) to determine that the GTRGAMMA substitution model was best. The GTRGAMMA model was run with 100 bootstrap replicates. We used SNAPP (through BEAST v.2.5; Bouckaert et al., 2019) to reconstruct a phylogeny that integrates across all possible gene trees to illustrate the group’s history using UCE data. We ran our SNAPP analysis for 30 million generations, sampling every 1000 steps, with a burn-in of 1 million generations. Tracer (v.1.7.2; Rambaut et al., 2018) was used to view the MCMC output to check for convergence and to ensure that no large-scale fluctuations were present in later trace trends. TreeSetAnalyzer (through SNAPP v.2.4.7; Bouckaert et al., 2019) was used to analyze the inferred gene trees to produce the 95% credible set of trees and to examine whether our mitochondrial topology was part of that set. SNAPP results were visualized using DensiTree (v.2.1.11; Bouckaert, 2010). For both our SNAPP and RAxML trees we included the mallard (*Anas platyrhynchos*) as an outgroup, giving us a confident root. For the mitogenomes we used MEGA to reconstruct a phylogeny using maximum likelihood with 100 bootstrap replicates. The mallard was again used as an outgroup. FigTree (v.1.4.2; Rambaut, 2006) was used to visualize resulting NEWICK files.

### 2.6. Demographic analysis

Diffusion Approximations for Demographic Inference (δaδi, v.1.7.0; Gutenkunst et al., 2009) was used to estimate demographic parameters under different models of pairwise divergence. Under a coalescent framework, δaδi predicts the joint frequency spectrum of genetic variation among populations enabling statistically rigorous assessments of user-defined demographic models focused on population size, gene flow rates, and divergence times (Gutenkunst et al., 2009). We tested eight models of divergence (Fig. S1): A) no divergence (neutral, populations never diverge); B) split with no migration (divergence without gene flow); C) split with migration (divergence with gene flow that is bidirectionally symmetric, 1 migration parameter); D) split with bidirectional migration (divergence with gene flow that is bidirectionally asymmetric, 2 migration parameters); E) split with exponential population growth, no migration; F) split with exponential population growth and migration; G) secondary contact with migration (1 migration parameter); and H) secondary contact with bidirectional migration (2 migration parameters). The scripts for these models are available here: https://figshare.com/s/e75158104c7896fa79e7. The neutral, split with migration, and exponential population growth models are provided in the δaδi file Demographics2D.py (as snm, splitmig, and IM, respectively). The models split-with-no-migration and split-with-exponential-growth-no-migration are versions of the split-with-migration (splitmig) and exponential population growth (IM) models with the migration parameters set to zero. The split-with-bidirectional-gene-flow model is a custom script that is a derivative of the split-with-migration (splitmig) model used to examine asymmetric gene flow. The secondary-contact model with one migration parameter (symmetric gene flow) is from Rougemont et al. (2017), and the secondary-contact model with two migration parameters is a derivative of that model to account for potential asymmetry in gene flow.

For each pairwise comparison, we ran a series of optimization runs, which consisted of running each model repeatedly under different parameters to find the most stable configuration with the lowest negative maximum log likelihood score (Table 2). Following this series of optimization runs, the best five log-likelihood scores from each set of subsequent runs were averaged to summarize that model, and we used the Akaike Information Criterion (AIC; Akaike, 1974; Burnham & Anderson, 2004) to determine the best-fit model. We then ran the best-fit model repeatedly and used demographic parameter estimates from this model’s top three runs to interpret values (as appropriate for each pairwise model) for ancestral population size (*Nref*, derived from *θ*), migration rates (*m* or *m12, m21*), current effective population sizes (*Ne* for each population, *nu1* and *nu2*), time since divergence (*T*), and time of secondary contact (*Tsc*), depending on model. The best-fit model was then bootstrapped to provide a 95% confidence interval around each parameter estimated (bootstrap_dadi.py v.1.1;https://github.com/jfmclaughlin92/beringia_scripts/).

To calculate substitution rates, we used BLASTn to compare our mallard reference FASTA to a fossil-calibrated node within the lineage of Anseriformes, the swan goose (*Anser cygnoides*; AnsCyg_PRJNA183603_v1.0) with a dated node of ∼28 Ma (Claramunt & Cracraft, 2015). Calculations for substitution rates, generation time, and adjusted length of sequences surveyed (Table S2) were then used with the best-fit model parameter estimates obtained from our demographic analyses to provide biological estimates of ancestral population size (*Nref*), size of populations (*nu1*, *nu2*), time since split (*T*), migration (gene flow in individuals/generation; derived from *m*), migration from population 1 into population 2 (individuals/generation; derived from *m12*), migration from population 2 into population 1 (individuals/generation; derived from *m21*), and time of secondary contact (*Tsc*) as appropriate (based on the best-fit model). The estimate for ancestral population size (Table S2) is derived from the output of *Θ* from δaδi; where *Θ =* 4*\**(*Nref*)*\**(substitution rate)*(adjusted length of sequences).

## 3. Results

### 3.1. Assembly and quality control

After assembly and filtering for UCE loci, we recovered 1,905 loci to create our *A. platyrhynchos* reference FASTA sequence. Following the bioinformatics pipeline for all individuals of read mapping, SNP calling, and quality filtering, the resulting dataset containing only high-quality data at all loci for all individuals (100% complete matrix) had 1,393 UCE loci, with a total length of 710,532 bp and a mean per-locus length of 510.1 bp (±3.9 bp, 95% CI).

The data matrix contained 1,204 variable loci and 189 invariable loci, and a total of 4,940 SNPs. For demographic analyses using δaδi, this high-quality dataset was thinned to be biallelic and contain 1 SNP/locus, which retained 1,202 variable loci. Identifying and removing Z-linked loci further reduced the number to 1,118 loci. Nucleotide diversity (π) was similar among the taxa (Table S3).

### 3.2. mtDNA and nuDNA phylogenies, and inferred gene tree history

The maximum likelihood phylogeny from the complete mitogenomes shows *A. c. carolinensis* sister to *A. flavirostris* and *A. c. crecca* sister to *A. c. nimia* (apart from a single *A. c. crecca* in the *carolinensis* clade, which is likely due to gene flow; Peters et al., 2012; Fig. 2A).

**Figure 2.**
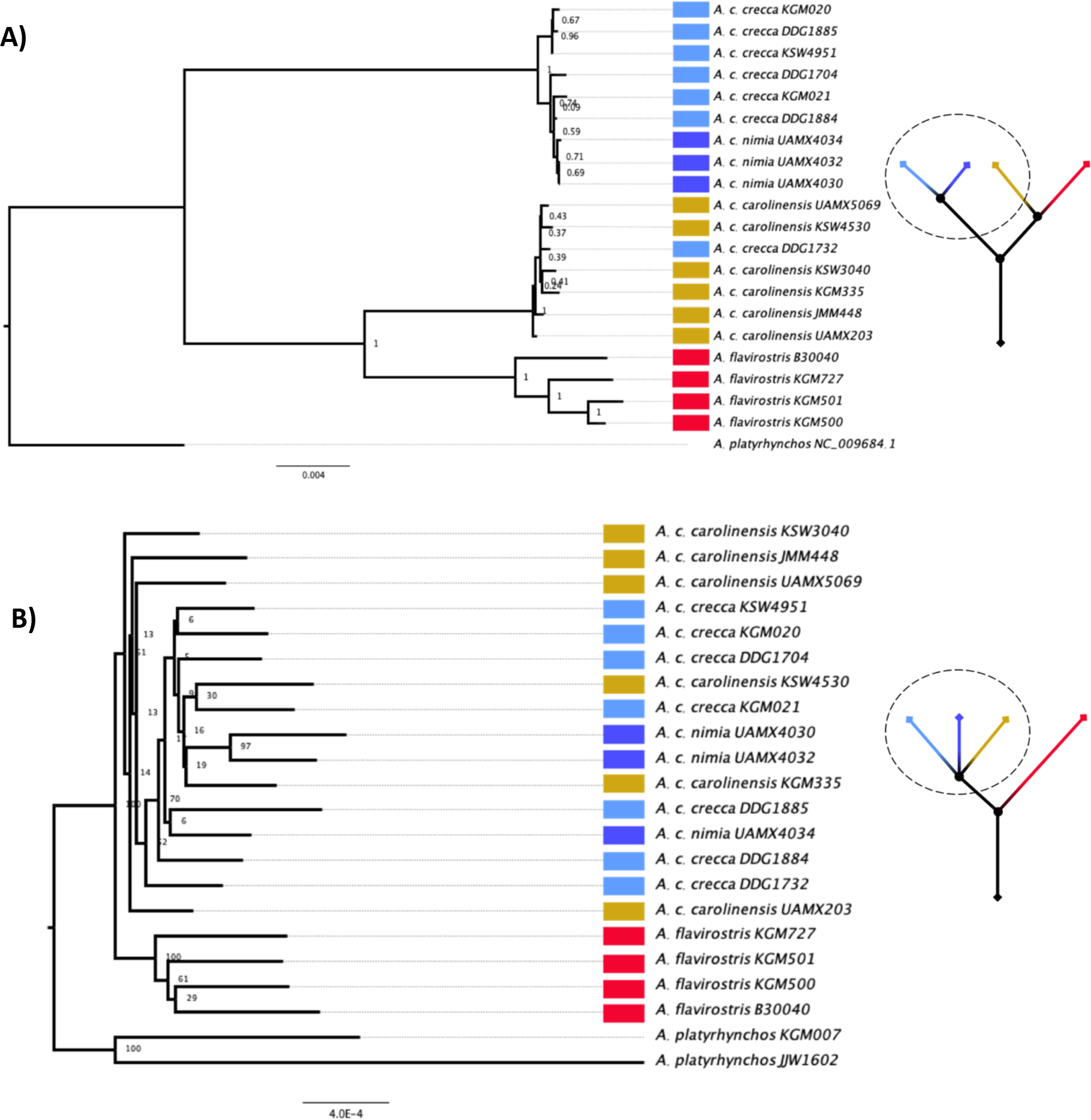
Maximum likelihood phylogenies of mitogenomic and nuclear DNA sequences. The mallard (*Anas platyrhynchos*) was used as an outgroup. Taxon colors correspond to Figure 1. **A)** Phylogeny of complete mitogenomes with 100 bootstrap replicates using MEGA (node values: 0-1). **B)** Phylogeny of UCEs with 100 bootstrap replicates (node values: 1-100), reconstructed using a 95% complete data matrix using RAxML. Values on internal nodes differ due to different programs being used for phylogenetic reconstruction. Adjacent to each phylogeny is a sketch illustration highlighting the discordance between the topologies, with the relationship of the *A. crecca* subspecies (within the circle) relative to the sister taxon *A. flavirostris*.

This topology is consistent with previous studies using mtDNA (Johnson & Sorenson, 1999, Gonzalez et al., 2009). Estimates of total mtDNA divergence were 3.7% between *A. c. crecca* and *A. c. carolinensis* and 2.2% between *A. c. carolinensis* and *A. flavirostris*. These divergences are commensurate with those reported in the literature (Johnson & Sorenson, 1999; Humphries & Winker, 2011) that indicate a deeper mtDNA split between *A. c. crecca* and *A. c. carolinensis* than between the latter and *A flavirostris*.

In contrast to mtDNA, which places *A. c. crecca* subspecies into two different well-resolved clades with one of these sister to *A. flavirostris* (Fig. 2A), our nuDNA (UCEs) phylogeny shows a much different topology, with all *A. crecca* individuals (*A. c. crecca, A. c. nimia*, and *A. c. carolinensis*) forming a polytomous clade that is sister to *A. flavirostris* (Fig. 2B). In this phylogeny, there was low bootstrap support for the nodes within *A. crecca*, resulting in a lack of structure at the subspecies level, whereas *A. flavirostris* individuals formed their own clade with 100% bootstrap support (Fig. 2B). The inferred UCE gene tree history from our SNAPP analysis showed the same sister relationship between *A. crecca* (*sensu lato*) and *A. flavirostris*, with considerable gene-tree conflicts evident among the *A. crecca* subspecies (Fig. 3). The mtDNA topology (Fig. 2A) thus falls outside of the 95% CI of the nuDNA SNAPP topology (Fig. 3), indicating that the two genomic histories are significantly different and reflecting mitochondrial-nuclear discord.

**Figure 3.**
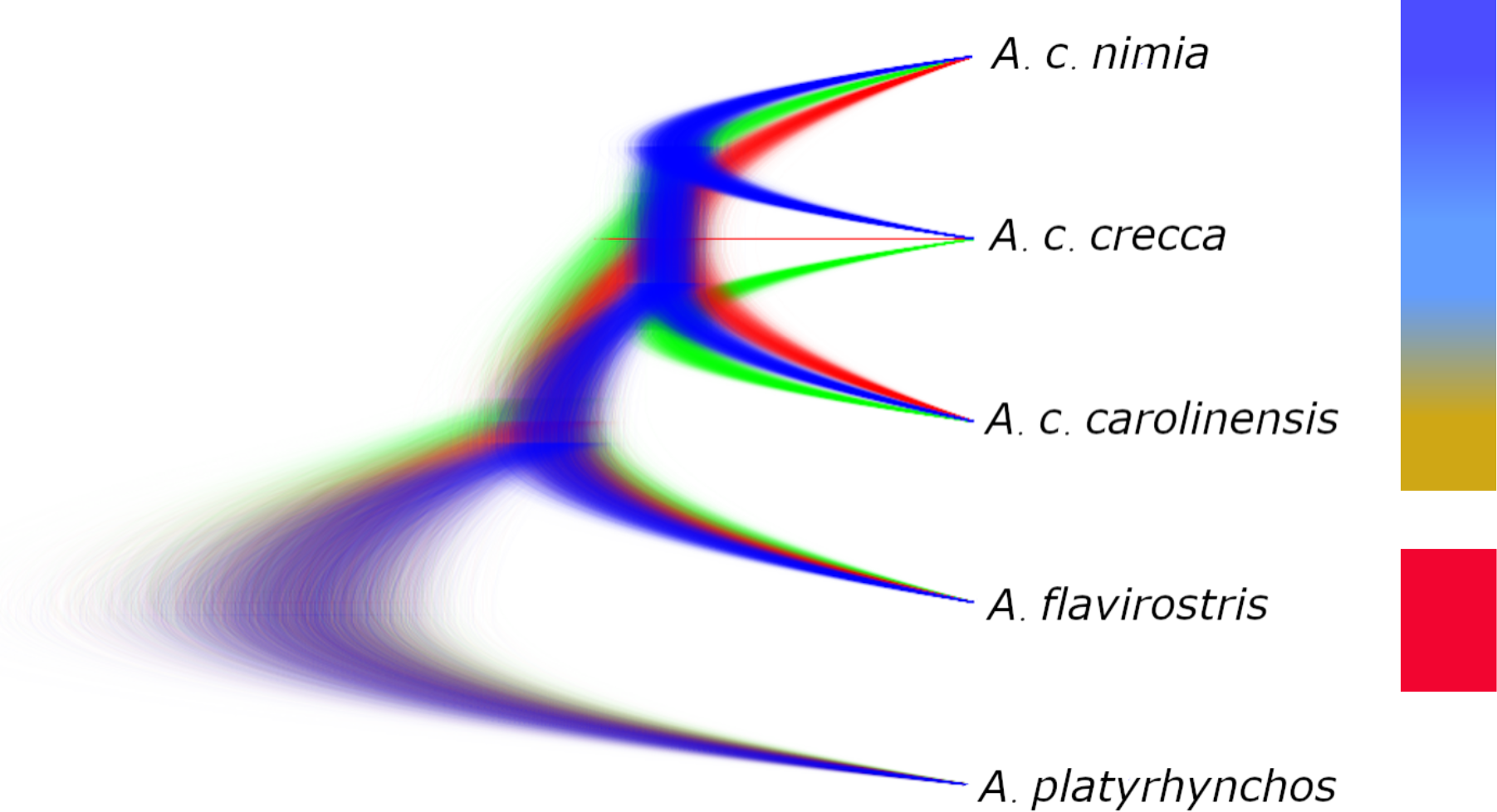
Inferred UCE gene tree history from a 30-million generation SNAPP analysis. Colors in the tree correspond to 1^st^- (blue), 2^nd^- (red), and 3^rd^-most (green) supported topologies. All individuals within the *A. crecca* complex form a polytomy and are illustrated by the colored gradient box, whereas the *A. flavirostris* clade is illustrated by the red box. The mallard (*A. platyrhynchos*) was used as an outgroup. Taxon colors correspond to Figure 1.

### 3.3. Divergence models and gene flow

*FST* values for each pairwise comparison (Table 1) ranged from a low of 0.033 (*A. c. crecca*−*A. c. carolinensis*; *P* = 0.03) to a high of 0.331 (*A. c. carolinensis*−*A. flavirostris*; *P* = 0.01). The *FST* value of 0.042 for *A. c. crecca*−*A. c. nimia* was not significant (*P* = 0.11), although higher than between *A. c. crecca*−*A. c. carolinensis* (Table 1). A principal components analysis (PCA) of *A. flavirostris* and the three subspecies of *A. crecca* using our thinned biallelic VCF file (Fig. 4) reflected the *FST* values, with the three *A. crecca* subspecies clustering closely together and *A. flavirostris* divergent from this group. The DAPC analysis also showed the *A. crecca* complex clustering together, with *A. flavirostris* being divergent (Fig. S2). One *A. c. crecca* individual clustered with *A. c. carolinensis* in our PCA plot (Fig. 4), possibly reflecting the well-known intercontinental hybridization between Holarctic subspecies in this system (Peters et al., 2012); the DAPC analysis clustered them all together (Fig. S2). The results for expected heterozygosity (*He*) versus observed heterozygosity (*Ho*) showed all comparisons to have significant differences, indicating a deviation from Hardy-Weinberg equilibrium and suggesting gene flow (Table S4).

**Table 1.** UCE-based population pairwise comparisons for between-population *FST* and their corresponding *P*-values. Asterisks (*) indicate significant differences.

| Pairwise Comparison | Between-population $F_{ST}$ | P-value ($F_{ST}$) |
| --- | --- | --- |
| *A. c. carolinensis*–*A. flavirostris* | 0.331 | 0.01* |
| *A. c. crecca*–*A. c. nimia* | 0.042 | 0.11 |
| *A. c. carolinensis*–*A. c. nimia* | 0.069 | 0.03* |
| *A. c. crecca*–*A. c. carolinensis* | 0.033 | 0.03* |

**Figure 4.**
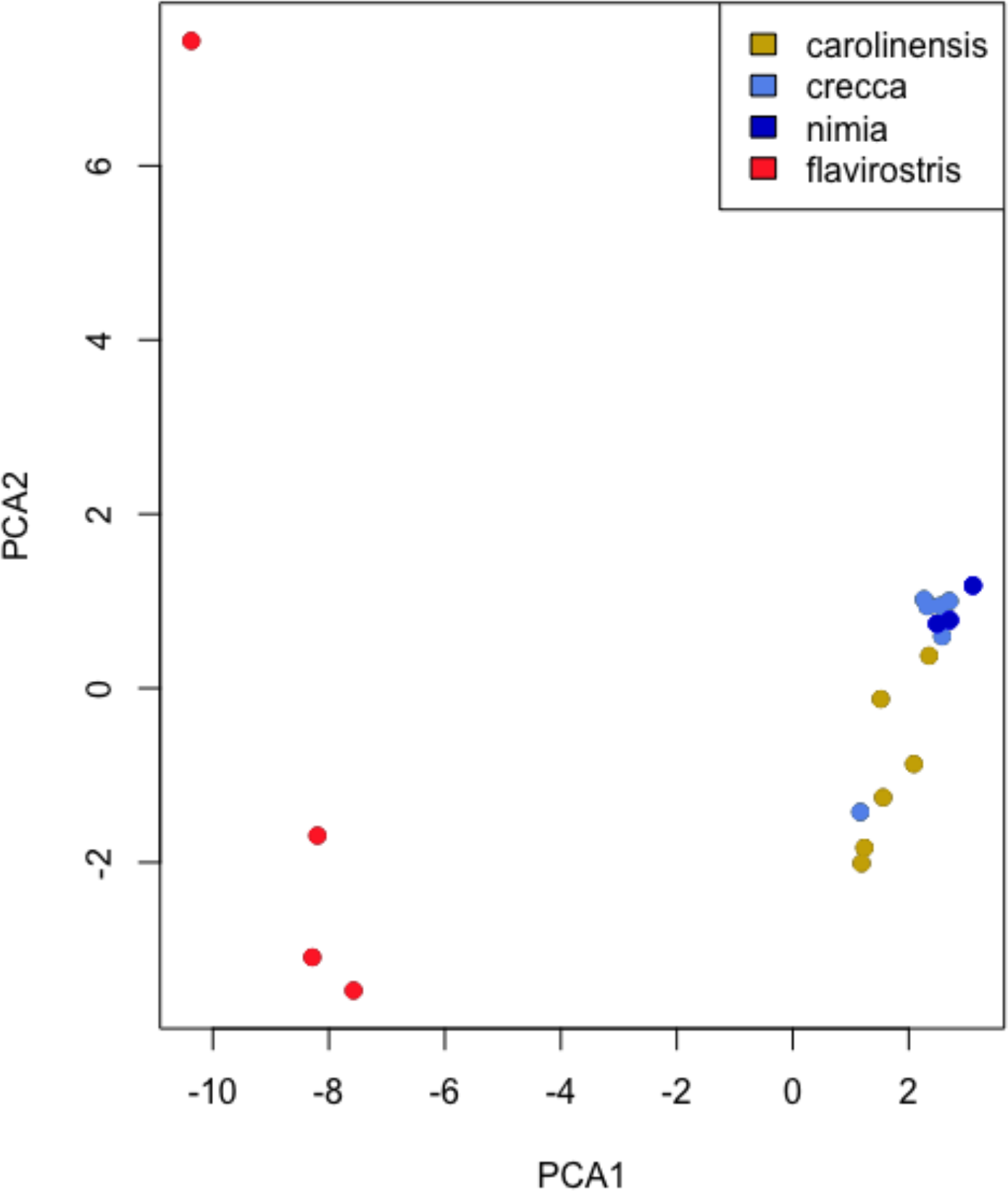
Principal components analysis (PCA) of *A. flavirostris* and all three subspecies of *A. crecca*. The one *A. flavirostris* outlier is the individual from Ecuador (Table S1), a different subspecies, *A. flavirostris andium*, likely indicating geographic isolation playing a role in the genetic variation found within *A. flavirostris* across its range in South America.

The best-fit models for our demographic analyses in δaδi found gene flow present in all pairwise comparisons (Fig. 5), as indicated by the AIC values (Table 2). When selecting best-fit models, some models had a ΔAIC of <10, which indicates models that are not statistically separable in their likelihood of explaining the data. Calculating ΔAIC causes the best model to have ΔAIC = 0, while the rest of the models have positive values (Burnham & Anderson, 2004). All of the statistically best-fit models included gene flow (Tables 2, S5). Among the *A. crecca* subspecies, there were effectively statistical ties between split-with-migration models (i.e., divergence with ongoing gene flow) and secondary-contact models (i.e., divergence with some isolation before resumption of gene flow; Tables 2, S5). For demographic analyses, we chose the best-fit model to be the one for each pairwise comparison with the lowest AIC value, a ΔAIC = 0, and a weighted AIC = 1 (Tables 2, S5). The best-fit models chosen were: split-with- symmetric-migration (Fig. S1C) for *A. c. carolinensis*−*A. c. nimia* and *A. c. crecca*−*A. c. nimia*; secondary-contact-with-bidirectionally-asymmetric-migration (Fig. S1H) for *A. c. crecca*−*A. c. carolinensis*; and split-with-symmetric-migration-and-exponential-population-growth (Fig. S1F) for *A. c. carolinensis*−*A. flavirostris*.

**Table 2.**
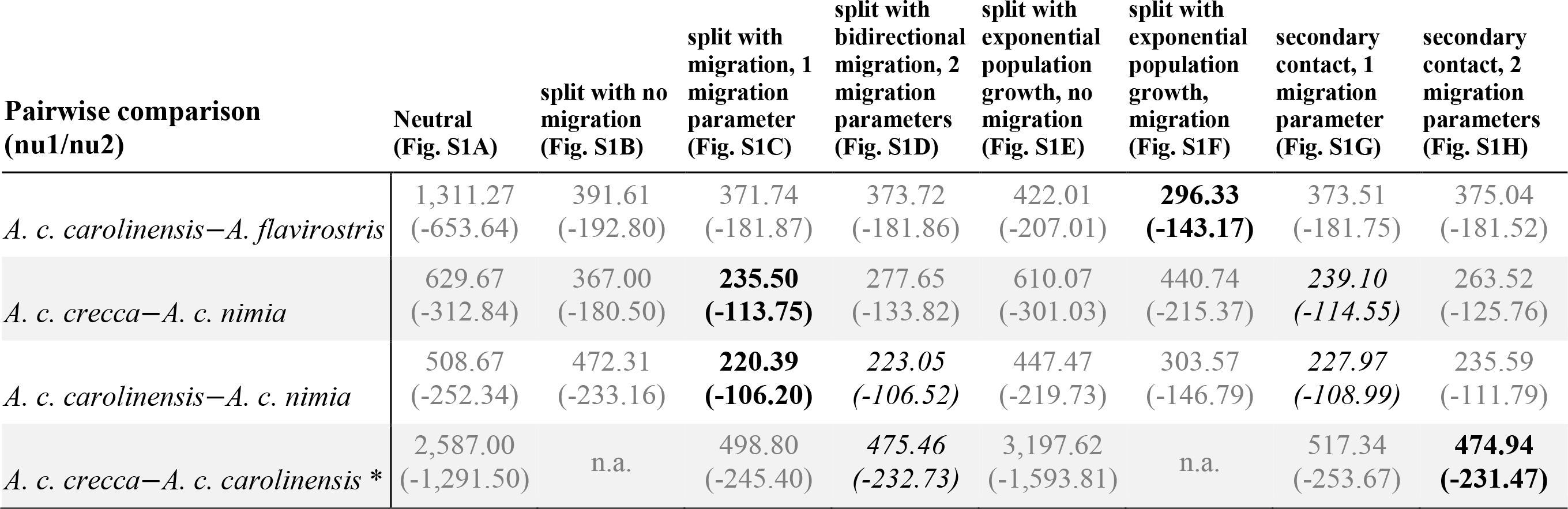
AIC values and negative log-likelihood values for each pairwise comparison for the eight demographic models tested (“migration” = gene flow). For each model, AIC values are paired with the negative log-likelihood values in parentheses. The negative log-likelihood values were averaged from the five best runs. Best-fit models with a weighted AIC of 1 are in bold, while runner-up models (ΔAIC < 10) are italicized. See Table S5 for ΔAIC values and weighted AIC values. *Results from McLaughlin et al. (2020). “n.a.” values indicate models that were unable to find a stable configuration and thus could not be run to completion (see McLaughlin et al. 2020, Table 3).

**Table 3.** Biological estimates obtained from the best-fit δaδi models for each pairwise comparison. Here we report ancestral population size (*Nref*, in number of individuals), size of population 1 (*Ne* or *nu1*, in number of individuals), size of population 2 (*Ne* or *nu2*, in number of individuals), migration (gene flow) from population 1 into population 2 (*m12* as individuals/generation), migration from population 2 into population 1 (*m21*, individuals/generation), time since split (*T*, in years), and time of secondary contact (*Tsc*, in years). See Figure 1.5 for a representation of the best-fit models. Values in parenthesis are the ±95% confidence interval around the biological estimates. *Results from McLaughlin et al. (2020).

| Pairwise comparison<br>(nu1/nu2) | Best fit model via<br>$\delta a \delta i$ | $N_{ref}$ | $N_e$ ($nu1$) | $N_e$ ($nu2$) | Migration<br>($m12$) | Migration<br>($m21$) | Time since<br>split ($T$) | Time of<br>secondary<br>contact<br>($T_{sc}$) |
| --- | --- | --- | --- | --- | --- | --- | --- | --- |
| *A. c. carolinensis*–*A. flavirostris* | split with migration &<br>exponential population<br>growth | 180,524<br>($\pm 9,731$) | 16,898,942<br>($\pm 231,289$) | 747,090<br>($\pm 59,819$) | 0.44<br>($\pm 0.09$) | 0.10<br>($\pm 0.04$) | 743,758<br>($\pm 38,436$) | - |
| *A. c. crecca*–*A. c. nimia* | split with migration | 138,010<br>($\pm 8,848$) | 4,369,087<br>($\pm 240,312$) | 530,970<br>($\pm 35,895$) | 8.78<br>($\pm 0.42$) | 1.07<br>($\pm 0.05$) | 561,742<br>($\pm 71,327$) | - |
| *A. c. carolinensis*–*A. c. nimia* | split with migration | 148,577<br>($\pm 11,955$) | 11,611,924<br>($\pm 210,055$) | 457,268<br>($\pm 64,171$) | 25.61<br>($\pm 1.08$) | 1.01<br>($\pm 0.04$) | 436,583<br>($\pm 113,537$) | - |
| *A. c. crecca* – *A. c. carolinensis* * | secondary contact &<br>bidirectional migration | 70,305<br>($\pm 1,967$) | 1,615,683<br>($\pm 39,884$) | 585,290<br>($\pm 38,181$) | 23.31<br>($\pm 0.63$) | 18.35<br>($\pm 2.29$) | 277,048<br>($\pm 12,993$) | 21,092<br>($\pm 10,979$) |

**Figure 5.**
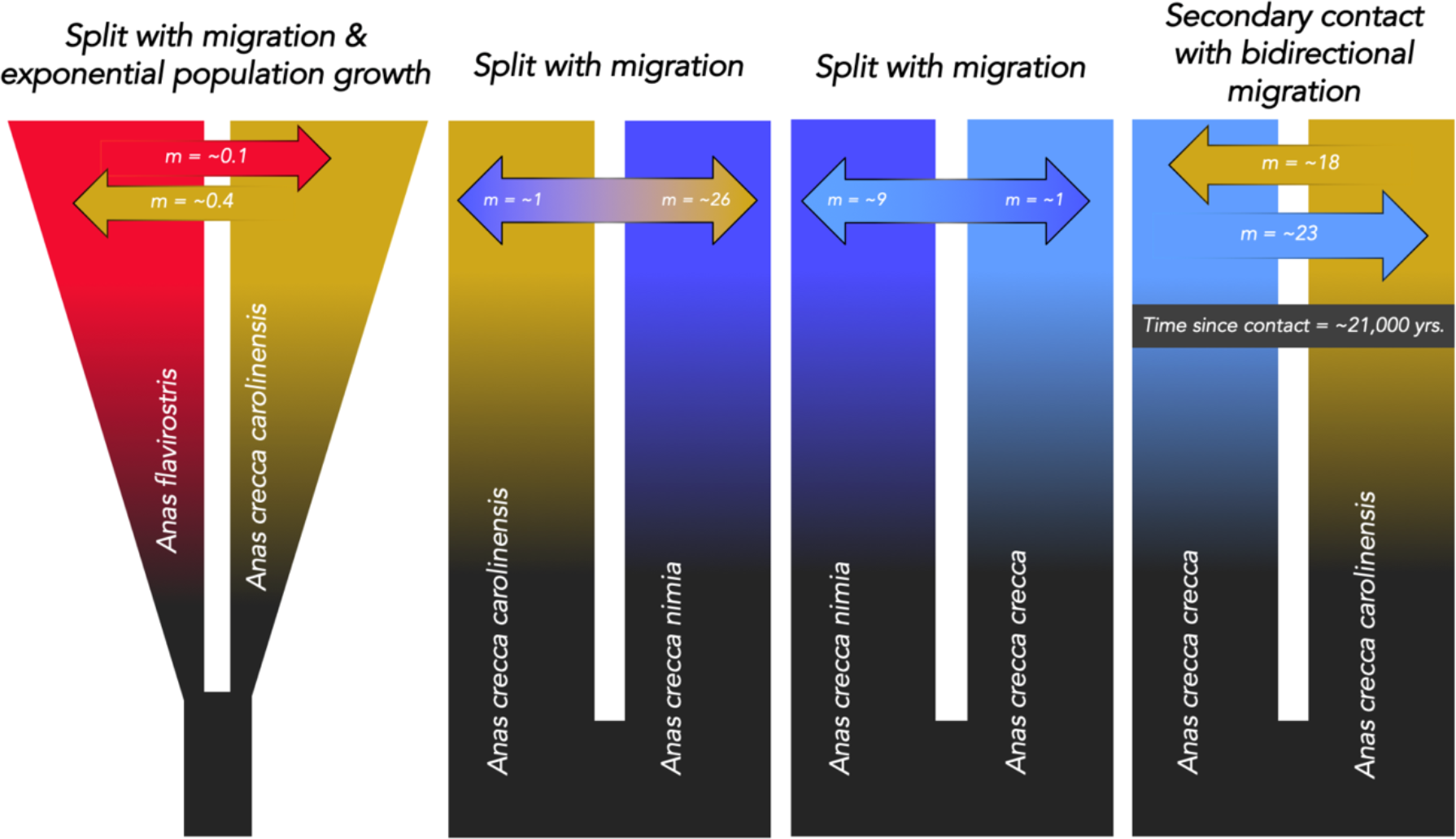
Visual representations of the best-fit demographic model for each pairwise comparison using δaδi (Gutenkunst et al., 2009). Migration (gene flow) rate estimates (*m*) are in individuals per generation. All analyses indicated divergence with gene flow. Technically, all fit in parapatric speciation theory, but, geographically, allopatry and heteropatry are involved. Full biological estimates are given in Table 3. Color coding corresponds to Figure 1. The *A. c. crecca−A. c. carolinensis* results are from McLaughlin et al., 2020.

The magnitude of inferred gene flow between these pairwise comparisons varied from ∼1 to ∼26 individuals per generation within the three *A. crecca* subspecies, with the highest levels of gene flow from *A. c. carolinensis* into *A. c. nimia* (Table 3, Fig. 5). The lowest levels of gene flow were ∼0.4 to ∼0.1 individuals per generation, which occurred in the *A. c. carolinensis*−*A. flavirostris* comparison (Table 3, Fig. 5). Estimates for ancestral population sizes (*Nref)* ranged from ∼70,300 individuals (in *A. c. crecca*−*A. c. carolinensis*) to 180,500 individuals (in *A. c. carolinensis*−*A. flavirostris*; Table 3). Values for effective population sizes (*Ne*) ranged from a low of ∼457,300 individuals (for *A. c. nimia* in *A. c. carolinensis*−*A. c. nimia*) to ∼16,900,000 individuals (for *A. c. carolinensis* in *A. c. carolinensis*−*A. flavirostris*; Table 3). UCE-based time since divergence (*T*) ranged from ∼277,000 years in *A. c. crecca*−*A. c. carolinensis* to ∼744,000 years in *A. c. carolinensis*−*A. flavirostris* (Tables 3, S6). The *A. c. crecca*−*A. c. carolinensis* comparison was the only one with the very best-fit model being one of secondary contact, in which time of secondary contact (*Tsc*) occurred ∼21,000 years ago coinciding with the end of the last glacial maximum (McLaughlin et al., 2020; Table 3, Fig. 5).

## 4. Discussion

Our study showed strong discordance between nuclear and mitochondrial phylogenies in the green-winged teal and yellow-billed teal complex (Figs. 2, 3). Our mitochondrial phylogeny using complete mitogenomes showed a sister relationship between *A. crecca carolinensis* and *A. flavirostris* and deeply divergent haplogroups within *A. crecca* (Fig. 2A), which agreed with previous, more limited mtDNA studies (Johnson & Sorenson, 1999; Gonzalez et al., 2009; Humphries & Winker, 2011; Peters et al., 2012; Peters et al., 2014). In contrast, nuclear phylogenies showed the *A. crecca* subspecies complex as an unresolved polytomy, with *A. flavirostris* as its sister (Fig. 2B, Fig. 3). Our other major finding is that divergence occurred with gene flow in all pairwise demographic analyses. We found no evidence of classic allopatric speciation, although levels of gene flow between *A. c. carolinensis* and *A. flavirostris* were low (less than one individual per generation), approaching the classic allopatric condition (as defined by Mayr, 1942; Gavrilets, 2003, 2004; Coyne & Orr, 2004; Fitzpatrick et al., 2008; Price, 2008). Periods of allopatry will foster divergence by allowing for reproductive isolation to develop.

However, given the absence of divergence models with secondary contact being supported as best-fit for the *A. c. carolinensis* and *A. flavirostris* comparison (Table 2), strict allopatric speciation (where the level of gene flow is zero) does not seem to be occurring.

### 4.1. Phylogenetic analyses of mtDNA and nuDNA reveal discord

There is strong evidence for mitonuclear discord in this group. The mitochondrial phylogeny showed *A. c. carolinensis* and *A. flavirostris* to be sister taxa, with *A. c. crecca* and *A*. *c. nimia* being sister to those (Fig. 2A). In contrast, our UCE phylogeny (Fig. 2B) and inferred UCE gene tree history (Fig. 3) showed *A. flavirostris* to form its own clade, sister to the *A. crecca* complex rather than to the subspecies *A. c. carolinensis* as indicated by the mitochondrial phylogeny (Fig. 2A). Furthermore, our mitogenomic phylogeny (Fig. 2A) matched previous mtDNA studies on this group (Johnson & Sorenson, 1999; Gonzalez et al., 2009). Després (2019) suggested that mitonuclear discord is a consequence of demographic fluctuations, for example when a large population is fragmented into small isolates, such as during glacial periods, which can result in founder events. Because mitochondrial DNA is more prone to lineage sorting because it has one-quarter the effective population size (*Ne*) of autosomal nuclear loci (Moore, 1995; Hudson & Turelli, 2003; Zink & Barrowclough, 2008), divergence can be stochastic with high potential to not reflect the underlying species tree. Given that Beringia has historically experienced cyclic fluctuations of climate change during the Pleistocene, climate fluctuations could have contributed to the mitonuclear discord within this group. Mitonuclear discord is prevalent in other Holarctic avian taxa that occur in Beringia, such as the mallard (*Anas platyrhynchos*; Peters et al., 2014), the Eurasian and American wigeons (*Mareca penelope – M. americana*; Humphries & Winker, 2011; Peters et al., 2014), and the *Eremophila* lark complex (Drovetski et al., 2014). However, unlike most birds, ducks have male-biased dispersal, and male-mediated intercontinental nuclear gene flow combined with female philopatry might be a leading contributor to this case of mtDNA and nuDNA discord (Peters et al., 2012, 2014; Fig. 3, Table 3).

We also note differences between the branch lengths in the mtDNA phylogeny (Fig. 2A) versus the nuDNA phylogeny (Fig. 2B). In the latter, the long terminal branch lengths could be due to recombination (Madison & Knowles, 2006; Kubatko et al., 2009; Lanier & Knowles, 2012). Recombination has not been effectively measured yet for UCE loci, but Winker et al. (2018) found loci exhibiting patterns indicative of recombination in around 15-25% of variable loci, suggesting that there are likely some recombinant haplotypes among our UCE sequences. Recombination within a locus could be a problem in sequence-based species tree reconstruction (Fig. 2B) because it might cause a mismatch between estimated gene trees and the true genealogies, and concatenation ignores recombination and makes the assumption that all loci have the same genealogy (Lanier & Knowles, 2012). However, the effects of recombination on species tree accuracy remain unclear and this has yet to be adequately addressed (Van Dam et al., 2021). Our inferred UCE gene tree history obtained through the SNAPP analysis (Fig. 3) uses one SNP per locus and therefore is not affected by recombination (although it might be affected by linkage disequilibrium; Bryant et al., 2012). In contrast, our UCE tree (Fig. 2B) provides a “winner take all” perspective through concatenation (Maddison, 1997), and while it does not account for recombination within a locus the majority of loci are likely not affected. Despite these two different methodologies, the two topologies are very similar, with the *A. crecca* complex forming a polytomous clade with *A. flavirostris* being sister to this complex.

Estimates of mitochondrial divergence show *A. c. crecca* and *A. c. carolinensis* to be more divergent (3.7%), than *A. c. carolinensis* and *A. flavirostris* (2.2%). Using smaller mtDNA datasets and substitution rates appropriate for those data (Peters et al., 2008; 2014), the dates of these divergences have been estimated at ∼2.6 Mya between *A. c. crecca*−*A. c. carolinensis* and ∼1.1 Mya between *A. c. carolinensis* and *A. flavirostris*. We note the differences here between estimates from mtDNA versus UCEs, with shallower splits found in UCEs (Table 3). Nuclear DNA generally has lower substitution rates than mitochondrial DNA, and so it might be concluded that the former are of less utility in estimating population divergence times (Arbogast et al., 2002). But male-mediated gene flow among *A. crecca* subspecies has also likely affected the nuclear *A. c. crecca—A. c. carolinensis* divergence time estimate, making it shallower. The relative depths of the nuclear splits (Table 1.3) also seem to be different than those depicted in the UCE SNAPP phylogeny (Fig. 1.3, blue, red, green), but we note that the pairwise comparisons with the youngest divergence dates also have the highest rates of gene flow inferred. Calibration points for estimating substitution rates also affect dating estimates (Arbogast et al., 2002), and might affect these mtDNA and nuDNA divergence estimates. If there had been no gene flow between Eurasian and North American populations of *A. crecca*, and substitution rate calibrations between marker types were accurate, we would expect to see the nuclear phylogeny match the mtDNA phylogeny, barring uncertainties stemming from incomplete lineage sorting. It has been established that deep mtDNA lineages do not always represent significant population divergences (Irwin, 2002; Zink & Barrowclough, 2008; Collins & Cruickshank, 2012; Morgan-Richards et al., 2017). This study highlights how biological species can have paraphyletic mitochondrial relationships, adding to many other cases of this taxonomically widespread phenomenon (Funk & Olmand, 2003).

The discordant phylogenies in these ducks make the biogeographic history uncertain. But lower female dispersal rates and male-mediated nuclear gene flow make it likely that the mtDNA history (Fig. 2A) is an accurate reflection of the biogeographic history of these lineages. The nuclear genome adds important information to our understanding of their subsequent divergence and speciation. Under this scenario, after Eurasia and North America were colonized during the Pleistocene (∼2.6 Mya) and mtDNA divergence between *A. c. crecca* and *A. c. carolinensis* was well established, South America was colonized by the proto-*carolinensis* ancestor of *A. flavirostris*. The intermittent or ongoing nuclear gene flow between Eurasian and North American populations confounded this biogeographic history, resulting in a single Holarctic biological species, while a later intercontinental colonist (*A. flavirostris*) emerged from this Holarctic complex, achieved a much higher degree of isolation, and became its own well-differentiated biological species that currently resides in South America and continues to differentiate (Fig. 4).

Modern *A. c. carolinensis* winter south almost to South America, and numerous records show individuals on the continent in Venezuela, Colombia, and Ecuador, and as far east as French Guiana (Meyer de Schauensee, 1966; Scott & Carbonell, 1986; Botero & Rusch, 1988; Renaudier et al., 2010; eBird, 2021). This wintering range might have been even farther south during prior glacial maxima (a period spanning ∼740 Kyr; Table 3). The phenomenon of migratory lineages dropping out new populations that establish new breeding grounds on or near tropical wintering grounds and subsequently differentiating and becoming new taxa occurs fairly often among numerous orders and families of birds (e.g., Bildstein, 2004; Winker, 2010; Winkler et al., 2017; Gómez-Bahamón et al., 2020). This seems to be a likely mechanism for the initial colonization of South America by the ancestors of *A. flavirostris*. It is also likely that a similar phenomenon, though at an individual level, caused the ongoing or intermittent gene flow between *A. c. carolinensis* and *A. flavirostris*. These birds form pair bonds on the wintering grounds, and it is plausible that wintering *A. c. carolinensis* males occasionally pair bond with *A. flavirostris* females and remain there to reproduce rather than return to northern breeding grounds. Our data suggest that over evolutionary time this occurs ∼4 times every ten generations, not inconceivable for a highly mobile source population with an effective population size of over 16 million birds (Table 3).

### 4.2. Divergence models reveal gene flow

We expected divergence with gene flow to occur among the three subspecies of *Anas crecca* given previous work, but our findings extend this understanding using a large number of orthologous nuclear loci and place the results in a directly comparable framework (Peters et al., 2012; Winker et al., 2013). We detected gene flow between the three subspecies of *A. crecca*, with relatively high levels between *A. c. crecca* and *A. c. carolinensis*, which have deeply diverged mtDNA lineages (Table 3, Fig. 2A). However, we could not statistically separate models of secondary contact versus ongoing gene flow for these subspecies (Table 2, S5). In addition, our *FST* results showing *A. c. crecca—A. c. nimia* to have an insignificant, but higher *FST* than the *A. c. crecca—A. c. carolinensis* comparison is counter to the findings of Winker et al. (2013). This latter result might be due to smaller sample sizes in our study. Optimal sample sizes for frequentist statistics (i.e., *FST, Ho, He*) are generally larger than optimal sample sizes for coalescent-based analyses (Felsenstein, 2005; Kalinowski, 2005; Morin et al., 2009; McLaughlin & Winker, 2020). Further work with more loci and perhaps larger sample sizes might provide better resolution among competing divergence-with-gene-flow models (Table 2) and for parameter estimates that are quite variable (e.g., effective population sizes; Table 3).

Interestingly our PCA analysis (Fig. 4) revealed that one *A. c. crecca* individual clustered within the *A. c. carolinensis* group, which was also reflected in our mitochondrial phylogeny (*A. c. crecca* DDG1732, Fig. 2A). This *A. c. crecca* individual likely reflects the hybridization known to occur in this system (Kulikova et al., 2005; Peters et al., 2007; Peters et al., 2012; Lavretsky et al., 2015).

### 4.3. Speciation modes

The pairwise comparisons within the *A. crecca* complex reaffirm the importance of parapatric and heteropatric (seasonally sympatric) divergence processes in this group, given the lineages’ geographic relationships and that gene flow was detected in all comparisons (Fig. 1.5; Table 1.2; Peters et al., 2012, Winker et al., 2013). Parapatric speciation, here occurring between Eurasian and North American continental lineages, is thought in this case to be driven by divergent selection stemming from sexual selection and different wintering grounds, although countered here by rather high levels of nuclear gene flow (Hartl & Clark, 1989; Rice & Hostert, 1993; Hostert, 1997; Price, 2008). However, this has resulted in only partial reproductive isolation between these populations, causing them to be stalled short of complete speciation, despite deeply divergent mitochondrial DNA (Peters et al., 2012). Heteropatric speciation is a type of ecological speciation driven by divergent selection occurring between lineages that are in sympatry and allopatry at different times during cyclic seasonal migrations (Winker, 2010; Winker et al., 2013). Both parapatric and heteropatric speciation involve gene flow between populations. However, periods of allopatric isolation (migration ≈ 0; Harrison, 2012), associated with glacial cycles at high latitudes and also seasonally at low latitudes, might have been important in lineage-specific evolutionary change, as seems possible in the *A. c. crecca*- *carolinensis* split and especially likely in *A. flavirostris* (Tables 2, 3). Finally, it is somewhat surprising that full allopatric isolation does not seem to have occurred in *A. flavirostris*, but that instead speciation with gene flow (albeit low) was strongly supported. The breeding ranges of *A. flavirostris* and *A. c. carolinensis* are fully allopatric and on different continents. But this isolation is apparently not complete, and strict allopatric divergence did not occur. This is probably the result of periodic recolonization of South America by migratory *A. c. carolinensis*, likely from their wintering grounds, which currently extend to northern Central America and the Northern Caribbean, with occasional birds recorded from South America.

### 4.4. Conclusions

We found mitonuclear discord within this group when comparing phylogenies reconstructed from the mitochondrial and nuclear genomes (Figs. 2, 3). Overall, our mitochondrial topology (Fig. 2A) matched relationships found in previous studies, in which *A. c. carolinensis* was sister to *A. flavirostris*, and *A. c. crecca* was sister to *A. c. nimia*. However, our nuclear phylogenies (Figs. 2B, 3) disagreed with the mitochondrial results, indicating that *A. crecca* subspecies formed one polytomy with *A. flavirostris* being sister to this group. This result could be explained by male-biased dispersal and the substantial levels of nuclear gene flow that we found among *A. crecca* subspecies (∼1–26 individuals per generation depending on pairwise comparison). We also found gene flow occurring between *A. c. carolinensis* and *A. flavirostris*, although these levels were low (∼0.1–0.4 individuals per generation). This suggests that while *A. flavirostris* is largely allopatric, *A. c. carolinensis* individuals might occasionally recolonize South America, for example through males pair-bonding in winter with a female *A. flavirostris* and remaining with her to reproduce rather than following a more species-appropriate appropriate mate back to her northern breeding grounds. We thus found no evidence for classic allopatric speciation (divergence without gene flow) in this group. Divergence with gene flow appears to be the predominant mode in this group, and the patterns of this gene flow have likely caused *A. crecca* (*sensu lato*) to be, mitochondrially, a paraphyletic biological species with respect to *A. flavirostris*.

## Acknowledgments

We thank the University of Alaska Museum and the Louisiana State University Museum of Natural Science for loans of tissue specimens and the specimen collectors who made it possible to do this research. We also thank the Kessel Fund for Northern Ornithology and the Friends of Ornithology for financial support, and Brant Faircloth for his input early in the project. This project was funded in part by the National Science Foundation [grant number DEB-1242267- 1242241-1242260], by an Institutional Development Award (IDeA) from the National Institute of General Medical Sciences of the National Institutes of Health [grant number 2P20GM103395] and by the National Institute of General Medical Sciences of the National Institutes of Health [grant number RL5GM118990]. Naoki Takebayashi and Devin Drown provided helpful comments and feedback on earlier drafts of the manuscript.

## Data Availability Statement

Original sequence data have been deposited in the NCBI Sequence Read Archive (SRA; Table S1; projects PRJNA741698 and PRJNA393740).

Additional data are available on figshare (https://figshare.com/s/f678d42ae94f520aef8e; https://figshare.com/s/4b22451c51ece33e4e60; https://figshare.com/s/245b3e7843b7c0622114): (a) the individual UCE sequences in a PHYLIP file (b) the reference UCE sequence data FASTA file, and (c) the VCF file used in δaδi analyses.

## Author Contributions

The study was conceived by Kevin Winker, Travis Glenn, Brant Faircloth, and Kevin McCracken. The data were generated by Travis Glenn and Brant Faircloth. The data were analyzed by Fern Spaulding. The first draft was written by Fern Spaulding. All authors contributed to the final version.

## Supplementary Information

**Table S1.** Specimen information of individuals used in this study. Note that UAM = University of Alaska Museum; LSUMZ = Louisiana State University Museum of Natural Science. All SRA reads available under project PRJNA741698 and project PRJNA393740. *Mallard individuals used as an outgroup. **Mallard individuals used as a reference.

| **Taxon** | **Locality** | **Tissue Number** | **Catalogue Number** | **Sequence Read Archive** |
| --- | --- | --- | --- | --- |
| *Anas crecca nimia* | Alaska: Aleutian Islands, Attu Island | UAM22887 | UAMX4032 | SRR15011467 |
| *Anas crecca nimia* | Alaska: Aleutian Islands, Attu Island | UAM22883 | UAMX4030 | SRR15011468 |
| *Anas crecca nimia* | Alaska: Aleutian Islands, Attu Island | UAM22877 | UAMX4034 | SRR15011476 |
| *Anas crecca carolinensis* | Alaska: Alaska Peninsula, Izembek NWR | UAM14961 | KGM335 | SRX2998946 |
| *Anas crecca carolinensis* | Alaska: Alaska Peninsula, Izembek NWR | UAM24497 | JMM448 | SRX2998977 |
| *Anas crecca carolinensis* | Alaska: Fairbanks | UAM11920 | KSW3040 | SRX2998949 |
| *Anas crecca carolinensis* | Alaska: Alaska Peninsula, Cold Bay | UAM20635 | KSW4530 | SRX2998947 |
| *Anas crecca carolinensis* | Alaska: Interior, Eielson Air Force Base | UAM11251 | UAMX203 | SRX2998948 |
| *Anas crecca carolinensis* | Alaska: Aleutian Islands, Chirikof Island | UAM28444 | UAMX5069 | SRX2998975 |
| *Anas crecca crecca* | Alaska: Aleutian Islands, Shemya Island | UAM9191 | DDG1732 | SRX2999024 |
| *Anas crecca crecca* | Alaska: Aleutian Islands, Shemya Island | UAM9255 | DDG1704 | SRX2998981 |
| *Anas crecca crecca* | Alaska: Aleutian Islands, Shemya Island | UAM14666 | DDG1884 | SRX2998973 |
| *Anas crecca crecca* | Alaska: Aleutian Islands, Shemya Island | UAM14100 | DDG1885 | SRX2998978 |
| *Anas crecca crecca* | Alaska: Aleutian Islands, Shemya Island | UAM11334 | KGM021 | SRX2998980 |
| *Anas crecca crecca* | Alaska: Aleutian Islands, Shemya Island | UAM11335 | KGM020 | SRX2998979 |
| *Anas crecca crecca* | Russia: Yakutia, Indigirka River | UAM22853 | KSW4951 | SRX2998972 |
| *Anas flavirostris oxyptera* | Bolivia: La Paz | UAM19017 | KGM500 | SRR15011474 |
| *Anas flavirostris oxyptera* | Bolivia: La Paz | UAM19018 | KGM501 | SRR15011473 |
| *Anas flavirostris flavirostris* | Argentina: Chubut | UAM19661 | KGM727 | SRR15011472 |
| *Anas flavirostris andium* | Ecuador: Napo Province | LSUMZ30040 | B30040 | SRR15011475 |
| *Anas platyrhynchos* * | Alaska: Interior, Chena River | UAM11193 | KGM007 | SRR15011471 |
| *Anas platyrhynchos* */** | Alaska: Interior, Eielson Air Force Base | UAM30286 | JJW1602 | SRR15011470 |
| *Anas platyrhynchos* ** | Alaska: Aleutian Islands, Shemya Island | UAM11328 | DDG1776 | SRR15011469 |

**Table S2.** Equations used to calculate biological estimates from δaδi output. Ancestral population size (*Nref*) is derived from the value of *Θ* obtained from δaδi. The equation for generation time (*G*) was obtained from Saether et al. (2005). For age of maturity *(⍺* ) we used a value of 1, given that most individuals attempt breeding in the first year and breed once every year thereafter (Bellrose, 1976). For annual adult survival *(s*), we used a value of 0.39, given that the mortality rates of adults (from banding recoveries) was 50–72% (Moisan et al., 1967; Johnson et al., 1992). The equations for adjusted base pair length (*adjusted L*) were obtained when calculating substitutions rates (Winker et al., 2018; Winker et al., 2019). To calculate substitution rates, we used a fossil-calibrated node within the lineage of Anseriformes, the swan goose (*Anser cygnoides*; NCBI: AnsCyg_PRJNA183603_v1.0) with a dated node of ∼28 Ma. (Claramunt & Cracraft, 2015).

| Biological Estimate | Equation |
| --- | --- |
| *generation time (G)*<br>*age of maturity ($\alpha$)*<br>*annual adult survival (s)* | $G = \alpha - [s/(1-s)]$ |
| *adjusted length (adjusted L)* | $\text{adjusted } L = (\text{length of sites surveyed}) * (\text{proportion of SNPs retained after thinning to 1 SNP/locus})$ |
| *Per site substitution rate (substitutions)* | $\text{substitutions} = \text{mutations/site/year}$ |
| *ancestral population size ($N_{ref}$)* | $N_{ref} = \Theta / [4 * (\text{substitutions/site/G}) * \text{adjusted } L]$ |

**Table S3.** Estimates of nucleotide diversity (π) from UCE data using concatenated data in MEGA (v.10; Kumar et al., 2018).

| **Taxon** | **Nucleotide Diversity ($\pi$)** |
| --- | --- |
| *A. crecca* subspecies and *A. flavirostris* | 0.000658 |
| *A. c. crecca* | 0.000583 |
| *A. c. carolinensis* | 0.000570 |
| *A. c. nimia* | 0.000531 |
| *A. flavirostris* | 0.000519 |

**Table S4.** UCE-based population pairwise comparisons for expected heterozygosity (*He*), observed heterozygosity (*Ho*), and their corresponding *P*-values. *P*-values for *He* vs. *Ho* were obtained from the Bartlett test of homogeneity of variances in adegenet (Jombart & Amhed, 2011). Asterisks (*) indicate significant differences.

| Pairwise Comparison | Expected<br>Heterozygosity ($H_e$) | Observed<br>Heterozygosity ($H_o$) | P-value<br>($H_e$ vs. $H_o$) |
| --- | --- | --- | --- |
| *A. c. carolinensis*–*A. flavirostris* | 0.0699 – 0.0691 | 0.0739 – 0.0578 | <2.2 x 10<sup>-16</sup>* |
| *A. c. crecca*–*A. c. nimia* | 0.0738 – 0.0592 | 0.0788 – 0.0729 | 2.87 x 10<sup>-05</sup>* |
| *A. c. carolinensis*–*A. c. nimia* | 0.0699 – 0.0592 | 0.0739 – 0.0729 | 0.0357* |
| *A. c. crecca*–*A. c. carolinensis* | 0.0738 – 0.0699 | 0.0788 – 0.0739 | 0.0411* |

**Table S5.** ΔAIC and weighted AIC values for each pairwise comparison for the eight demographic models tested (“migration” = gene flow). For each model, the ΔAIC are listed with the weighted AIC values in parentheses. Best-fit models with a ΔAIC of 0 and weighted AIC of 1 are in bold. Models that are not statistically separable from the best-fit models (ΔAIC of <10) are italicized. *Results from McLaughlin et al. (2020). “n.a.” values indicate models that were unable to find a stable configuration and thus could not be run to completion (see McLaughlin et al. 2020, Table 3).

| Pairwise comparison<br>(nu1/nu2) | Neutral<br>(Fig. S1A) | split with no<br>migration<br>(Fig. S1B) | split with<br>migration, 1<br>migration<br>parameter<br>(Fig. S1C) | split with<br>bidirectional<br>migration, 2<br>migration<br>parameters<br>(Fig. S1D) | split with<br>exponential<br>population<br>growth, no<br>migration<br>(Fig. S1E) | split with<br>exponential<br>population<br>growth,<br>migration<br>(Fig. S1F) | secondary<br>contact, 1<br>migration<br>parameter<br>(Fig. S1G) | secondary<br>contact, 2<br>migration<br>parameters<br>(Fig. S1H) |
| --- | --- | --- | --- | --- | --- | --- | --- | --- |
| *A. c. carolinensis*—*A. flavirostris* | 1,014.94<br>(4.06x10<sup>-221</sup>) | 95.27<br>(2.05x10<sup>-21</sup>) | 75.41<br>(4.22x10<sup>-17</sup>) | 77.38<br>(1.57x10<sup>-17</sup>) | 125.68<br>(5.12x10<sup>-28</sup>) | **0.00**<br>**(1.00)** | 77.17<br>(1.75x10<sup>-17</sup>) | 78.71<br>(8.10x10<sup>-18</sup>) |
| *A. c. crecca*—*A. c. nimia* | 394.18<br>(2.54x10<sup>-86</sup>) | 131.51<br>(2.78x10<sup>-29</sup>) | **0.00**<br>**(1.00)** | 42.15<br>(7.03x10<sup>-10</sup>) | 374.57<br>(4.61x10<sup>-82</sup>) | 205.25<br>(2.70x10<sup>-45</sup>) | *3.60*<br>*(0.17)* | 28.02<br>(8.21x10<sup>-7</sup>) |
| *A. c. carolinensis*—*A. c. nimia* | 288.28<br>(2.52x10<sup>-63</sup>) | 251.92<br>(1.98x10<sup>-55</sup>) | **0.00**<br>**(1.00)** | 2.65<br>*(0.27)* | 227.08<br>(4.91x10<sup>-50</sup>) | 83.18<br>(8.66x10<sup>-19</sup>) | 7.58<br>*(0.02)* | 15.19<br>(5.02x10<sup>-4</sup>) |
| *A. c. crecca*—*A. c. carolinensis* * | 2,112.06<br>(0.00) | n.a. | 23.86<br>(6.59x10<sup>-6</sup>) | 0.52<br>*(0.77)* | 2720.68<br>(0.00) | n.a. | 42.40<br>(6.21x10<sup>-10</sup>) | **0.00**<br>**(1.00)** |

**Table S6.**
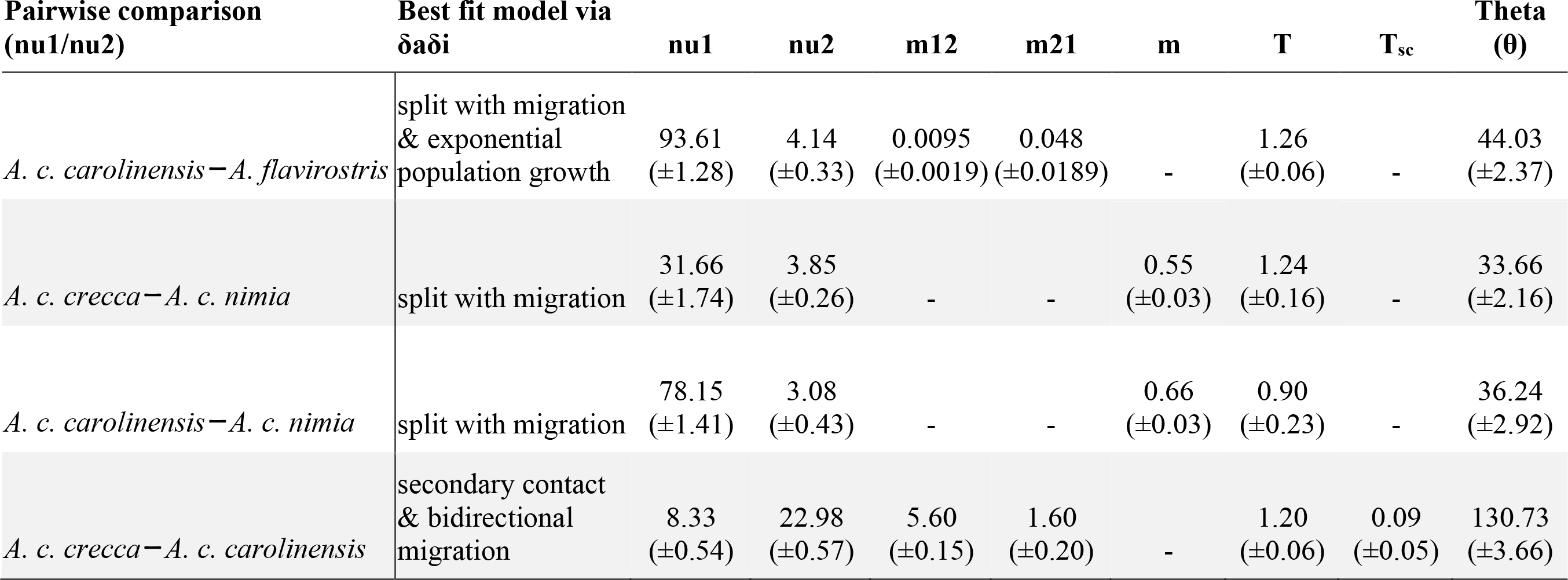
Raw parameter output from δaδi used to calculate biological estimates. The average of the top three runs output from the δaδi best-fit model for each pairwise comparison is paired with the ±95% confidence interval around that parameter (in parentheses). Instances of a hyphen indicated that the parameter is not present in that best-fit model. These values were translated into biological estimates using the equations listed in Table S2. Translated values are listed in Table 3.

**Figure S1.**
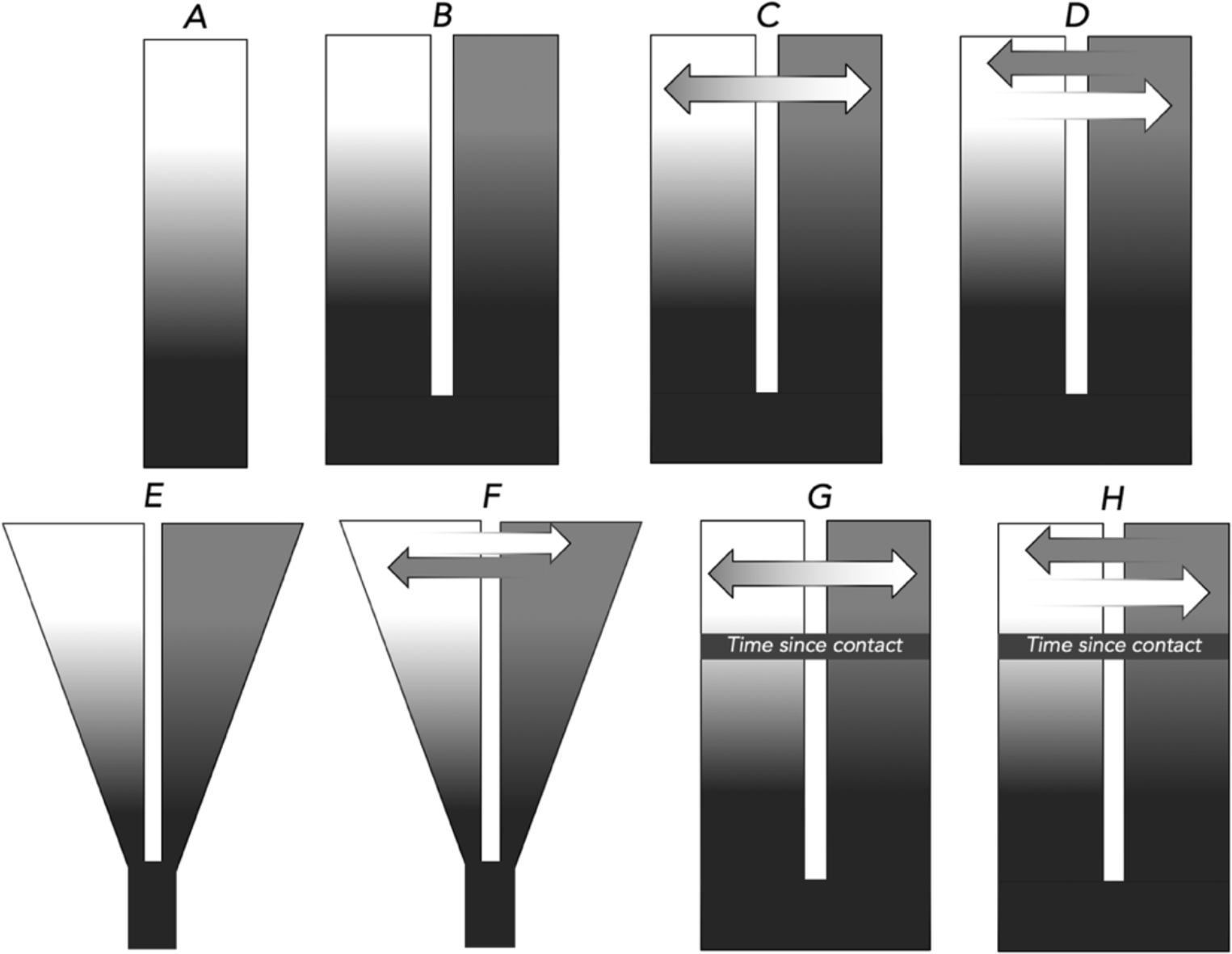
Models of divergence and speciation tested using δaδi (Gutenkunst et al., 2009) to determine demographic histories between populations: **A)** neutral, no divergence; **B)** split with no migration; **C)** split with migration, 1 migration parameter (i.e., bidirectionally symmetric); **D)** split with bidirectional migration, 2 migration parameters (i.e., bidirectionally asymmetric); **E)** split with exponential population growth, no migration; **F)** split with exponential population growth and migration; **G)** secondary contact with migration (1 migration parameter); and **H)** secondary contact with bidirectional migration (2 migration parameters). Models that contain one arrow indicate gene flow at relatively equal levels, while models with two arrows indicate unequal levels of gene flow (asymmetric). The gradient of contrast illustrates increasing population differentiation.

**Figure S2.**
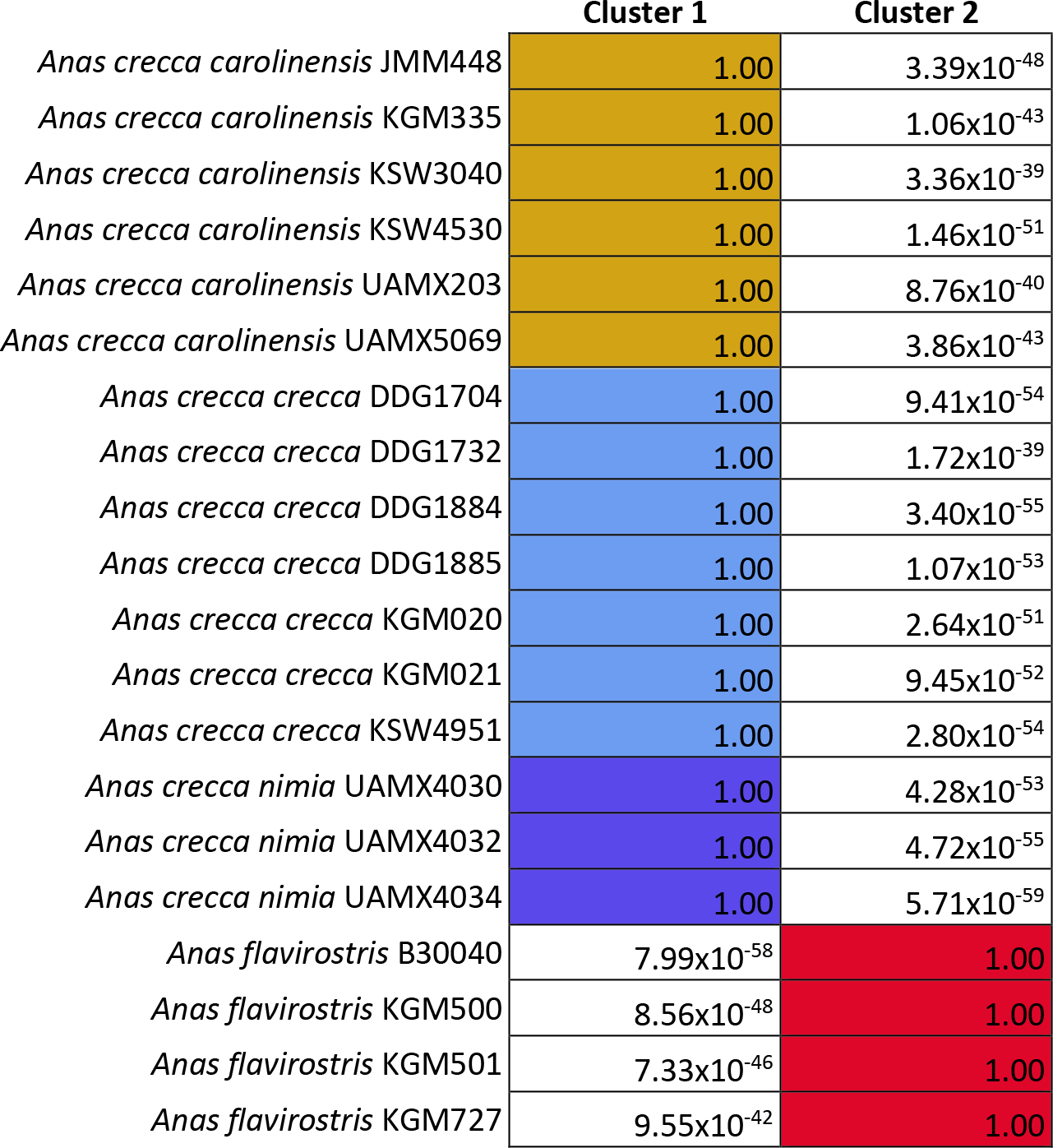
DAPC cluster assignment plot of the three *A. crecca* subspecies and *A. flavirostris*, with 20 PCAs retained and 2 clusters. DAPC posterior values retained for DAPC cluster assignment plot of the three *A. crecca* subspecies and *A. flavirostris*, with a prior group assignment of 2 clusters. Each individual contains the species name and field catalogue number. Colors correspond to Figure 1.

## Notes

### Competing Interest Statement

The authors have declared no competing interest.

